# RanBP1 plays an essential role in directed migration of neural crest cells during development

**DOI:** 10.1101/2022.05.05.490747

**Authors:** Elias H Barriga, Delan N Alasaadi, Chiara Mencarelli, Roberto Mayor, Franck Pichaud

**Author notes:** Co-first authorship (Elias H Barriga, Delan N Alasaadi).

## Abstract

Collective cell migration is essential for embryonic development, tissue regeneration and repair, and has been implicated in pathological conditions such as cancer metastasis. It is, in part, directed by external cues that promote front-to-rear polarity in individual cells. However, our understanding of the pathways that underpin the directional movement of cells in response to external cues remains incomplete. To examine this issue we made use of neural crest cells (NC), which migrate as a collective during development to generate vital structures including bones and cartilage. Using a candidate approach, we found an essential role for Ran-binding protein 1 (RanBP1), a key effector of the nucleocytoplasmic transport pathway, in enabling directed migration of these cells. Our results indicate that RanBP1 is required for establishing front-to-rear polarity, so that NCs are able to chemotax. Moreover, our work suggests that RanBP1 function in chemotaxis involves the polarity kinase LKB1/PAR4. We envisage that regulated nuclear export of LKB1 through Ran/RanBP1 is a key regulatory step required for establishing front-to-rear polarity and thus chemotaxis, during NC collective migration.

## INTRODUCTION

Polarity is a widespread feature in biology, and the function of many cell types is rooted in their ability to become polarised. This is the case for neurons, with the soma-axon polarity axis, for epithelial cells, with the apical (top) – basal (bottom) axis, and for migrating cells, which can polarise along a front-to-rear axis. While progress continue to be made on our understanding of the pathways that promote cell polarity, gaps remain, especially considering how information in a cell’s microenvironment can mobilise cellular machineries to generate or remodel polarity. In previous work using a genetic approach to address this importat issue in biology, we found that the Ran pathways, which controls nucleocytoplasmic exchange of macromolecules, is required for radial migration and polarisation of cortical neurons (i.e. axon specification), during cortex development (Mencarelli et al., 2018). During nuclear export, RanBP1 promotes cargoes unloading from the Ran-export complex, which terminates cargo export and allows for the import of Ran, back to the nucleus (Bischoff and Gorlich, 1997). Remarkably, failure in differentiating an axon in cortical neurons deficient for RanBP1 could be rescued by expressing the polarity factor, serine/threonine kinase, LKB1/PAR4 (Mencarelli et al., 2018), a known Ran cargo (Baas et al., 2003; Dorfman and Macara, 2008; Matsuki et al., 2010). This led us to propose that regulated nuclear export of LKB1 is a key step in establishing polarity in cortical neurons, a notion well supported by other studies in these cells and also in epithelial cells (Baas et al., 2004; Barnes and Polleux, 2009; Matsuki et al., 2010; Shelly et al., 2007). Whether a similar RanBP1-LKB1 connection plays a role in directed migration has not been explore so far.

Next to cortical neurons, RanBP1 is also highly transcribed in the eye and in the cranial neural crest cells (NC) (Maynard et al., 2003). In addition, velo-cardio-facial/DiGeorge syndrome, a developmental disease caused by a micro-deletion (22q11.2) uncovering RanBP1 (Dunham et al., 1999; Merscher et al., 2001), includes developmental defects that can be attributed to the NC cells (McDonald-McGinn et al., 2015). Furthermore, NC migration requires LKB1 (Creuzet et al., 2016). These observations prompted us to examine whether the RanBP1-LKB1 pathway plays a role in NC migration. Early on in vertebrate development, the NCs migrate as a collective, and populate the embryo in a stereotyped manner to induce formation on many adult structures and organs, including bones, cartilage, and portions of the heart, for example (Etchevers et al., 2019; Szabo and Mayor, 2018). Collective cell migration can be defined as a cooperative mode of cellular motion whereby group of cells migrate in a highly coordinated manner. This mode of migration is vital for a range of biological processes during embryogenesis, wound healing and cancer metastasis (Barriga and Mayor, 2019; Mayor and Etienne-Manneville, 2016; Rorth, 2009, 2011). During collective cell migration, the direction of migration is set though biochemical or biophysical cues, which are present in the microenvironment (Barriga and Theveneau, 2020; Rorth, 2009; Shellard and Mayor, 2020). These cues direct migration by activating signalling downstream of receptors, which in turn can harness the activity of cell intrinsic pathways to promote directed cell motility.

Here, we examine the function of the RanBP1-LKB1 pathway in NC migration. We find that RanBP1 is required for NC migration *in vivo,* and our results indicate that this is in part due to a function in enabling chemotaxis. Furthermore, our results show that while LKB1 knockdown affects NC migration, overexpression of this kinase can rescue the migration defect induced by the knockdown of RanBP1. Altogether, our results suggest that RanBP1 requirment in NC migration depends upon the polarity effector LKB1. Our work supports the idea that regulated nucleocytoplasmic transport of this kinases plays an important role in cell across cell types.

## RESULTS

### RanBP1 is required for neural crest cell migration *in vivo*

In order to test RanBP1 function in *Xenopus* NC cells *in vivo,* we microinjected a morpholino (MO) specifically designed to inhibit RanBP1 expression (RanBP1-MO). The effect of this morpholino on NC cells migration was analysed by monitoring NC position using *in situ* hybridisation (ISH) against *snail2,* a marker for migrating NC. In these experiments, one side was injected with the MO and the pattern of NC was compared to that of the contralateral control side of the same embryo. We determined that injecting 6.5 ng of RanBP1-MO *per* blastomere was an optimal concentration whereby we could achieve a strong inhibition of RanBP1 as shown by western blot analysis (Figure 1A-B), without affecting embryo survival.

**Figure 1:**
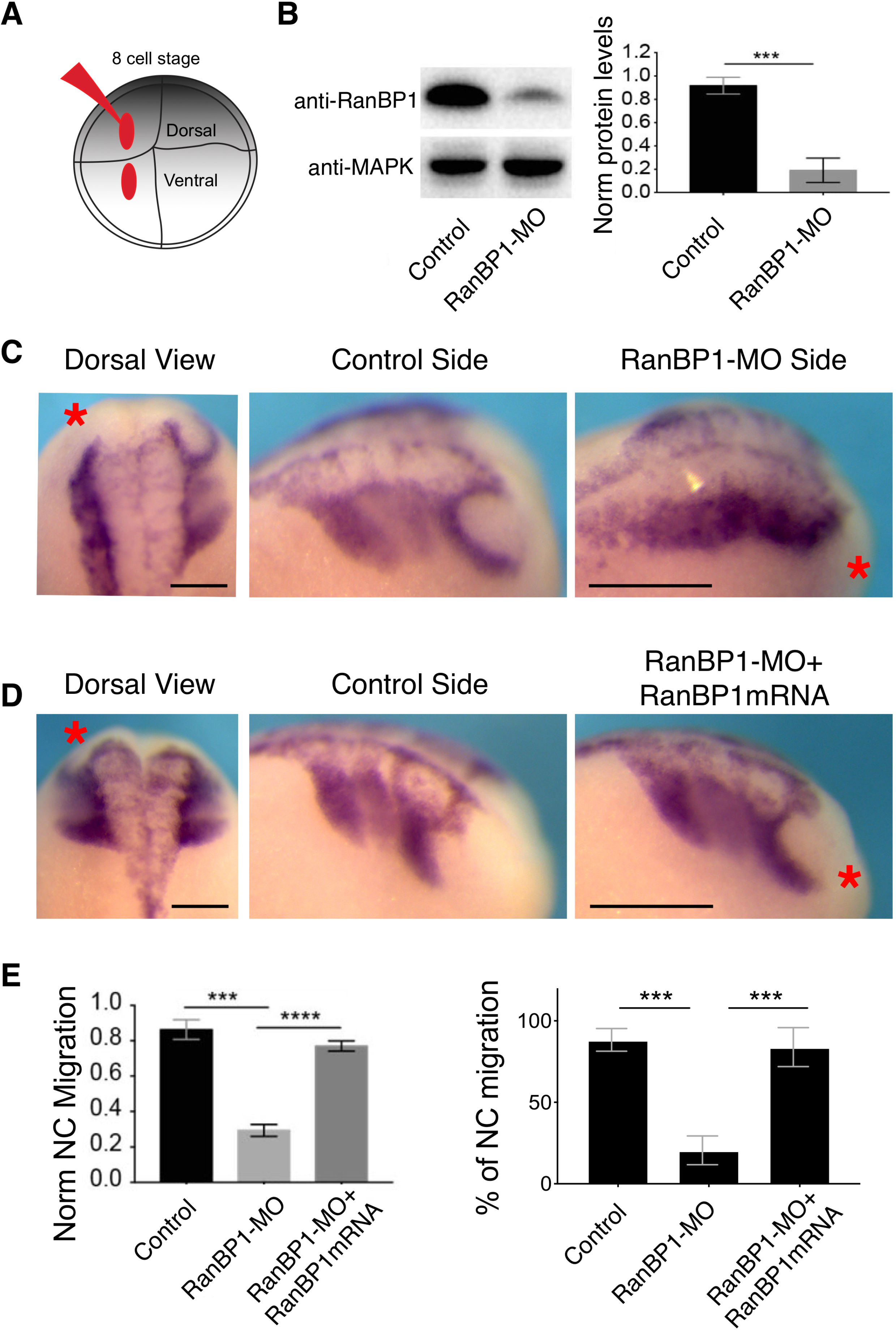
RanBP1 is required for neural crest migration. (**A**) Diagram depicting the microinjection strategy. (**B**) Western blot of cephalic *Xenopus* regions enriched in NCs from stage 21-22 animals, and gel densitometry showing RanBP1 levels in Control and RanBP1-MO embryos; histogram represents the media and bars the standard deviation of three independent experiments, two-tailed t-test ***, P < 0.001, N = 3 blots from 3 independent experiments. (**C** and **D**) Dorsal and lateral views of embryos hybridized with a probe against *snail2* to label migrating NCs. (**C**) a red asterisk marks the injected side of the embryo. Lateral views show that while NCs from the control side of the embryo migrate normally by forming dorso ventral streams, RanBP1-MO cells (red asterisk) stay in the dorsal part of the embryo. (**D**) a red asterisk marks the injected side of the embryo and dorsal as well as lateral views show that both Control and RanBP1-MO+RanBP1mRNA co-injected NCs migrate normally by forming dorso ventral streams. Scale bars represent 200 μm. (**E**) Normalized NC migration and percentage of embryos displaying the representative phenotypes shown in (**C** and **D**). Histograms represent the median and bars the standard deviation of three independent experiments, N= 20 embryos for each condition. two-tailed t-test ***, P < 0.001 and ****, P < 0.0001.

While NCs normally migrate by forming stereotypical streams from dorsal to ventral territories, we found that RanBP1-MO treated cephalic NC cells failed to migrate (Figure 1C-E). Instead, they remained as a group at the border of the neural folds, which is the region of the embryo where they are originally induced (Figure 1C and 1E). Rescue experiments using a human RanBP1 (hRanBP1) mRNA (Mencarelli et al., 2018), which cannot be targeted by the MO, confirmed that this phenotype was due to the specific loss of RanBP1 (Figure 1D-E). To ensure that the MO RanBP1 migration phenotype was not due to defects in sanil expression, we made use of other, complementary migratory NC makers, such as *FoxD3, Twist* and *Snail2.* These markers confirmed that NC migration was impaired when decreasing the expression of RanBP1 (Figure 2). In order to test wether RanBP1 depletion could have an effect on early NC specification or ectodermal development in general, we analyzed the expression of Pax3, Sox2 and Keratin. This analysis showed that RanBP1-MO does not affect the expression of early neural fold (Figure 3A,G) or early NC (Figure 3B-D,G) markers. Morevover, RanBP1-MO did not affect the expression of epidermal or neural plate markers (Figure 3E-G). To further characterize the RanBP1-MO phenotype in *Xenopus,* we also examined the effect of RanBP1 loss-of-function in craniofacial development, a well-known NC derivate tissue (Barriga et al., 2013). For this purpose, stage 46 larvae were stained with alcian blue to specifically visualise the cartilage. This revealed multiple defects in cranial cartilage morphology on the MO-injected side of the embryo when compared to the non-treated one (Figure 4). Altogether, these results are consistent with a requirement for RanBP1 in NC migration during development.

**Figure 2:**
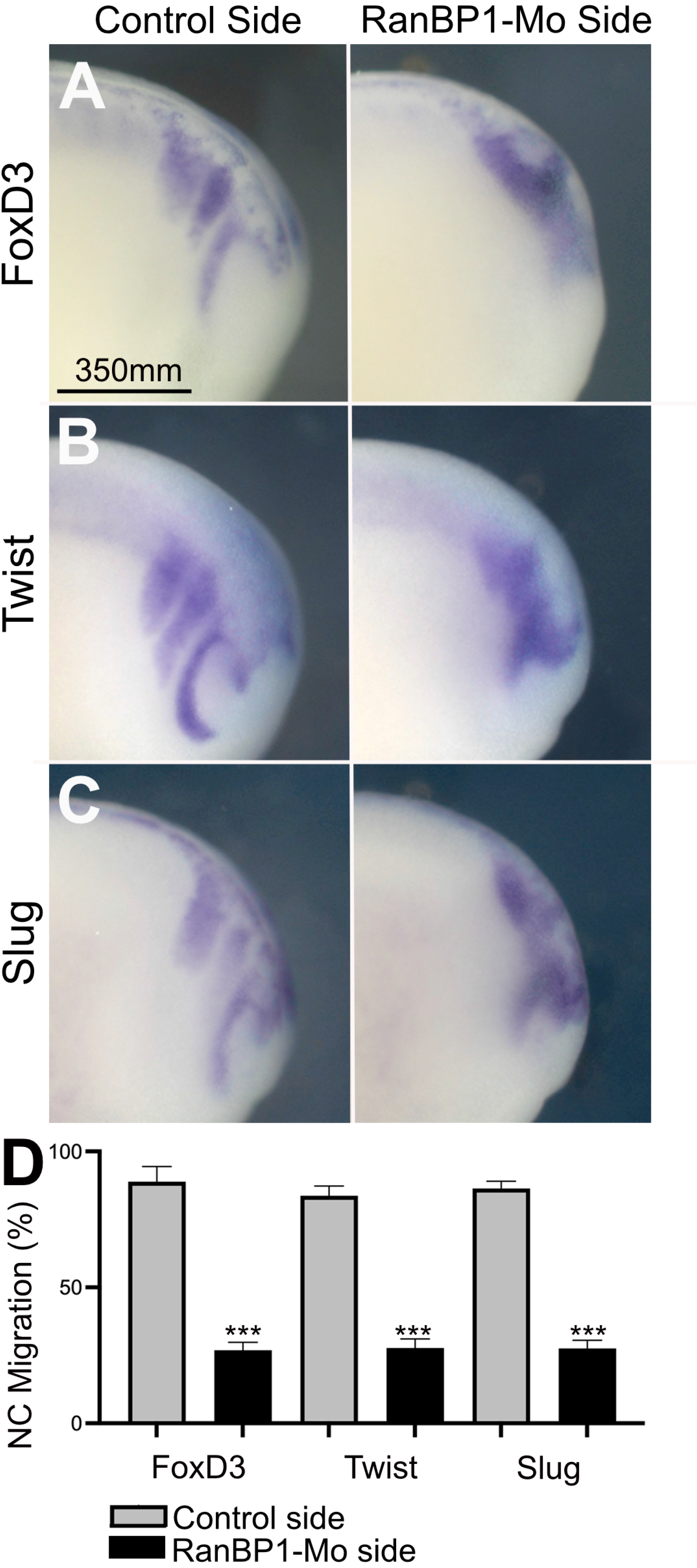
Analysis of NC migration. Embryos were injected as described in Figure 1, and NC migration was analyzed by ISH against FoxD3 in stage 24 animals (**A**), Twist (**B**) and Slug (**C**). Lateral views of the same embryo show the uninjected control side and injected side, as indicated. Inhibition of NC migration is observed in stage 24 animals for the 3 markers upon RanBP1 depletion. (**D**) Percentage of stage 24 embryos displaying normal NC migration. Histograms represent the median and bars the standard deviation of three independent experiments, N= 15 stage 24 embryos for each condition. two-tailed t-test ***, P < 0.001.

**Figure 3:**
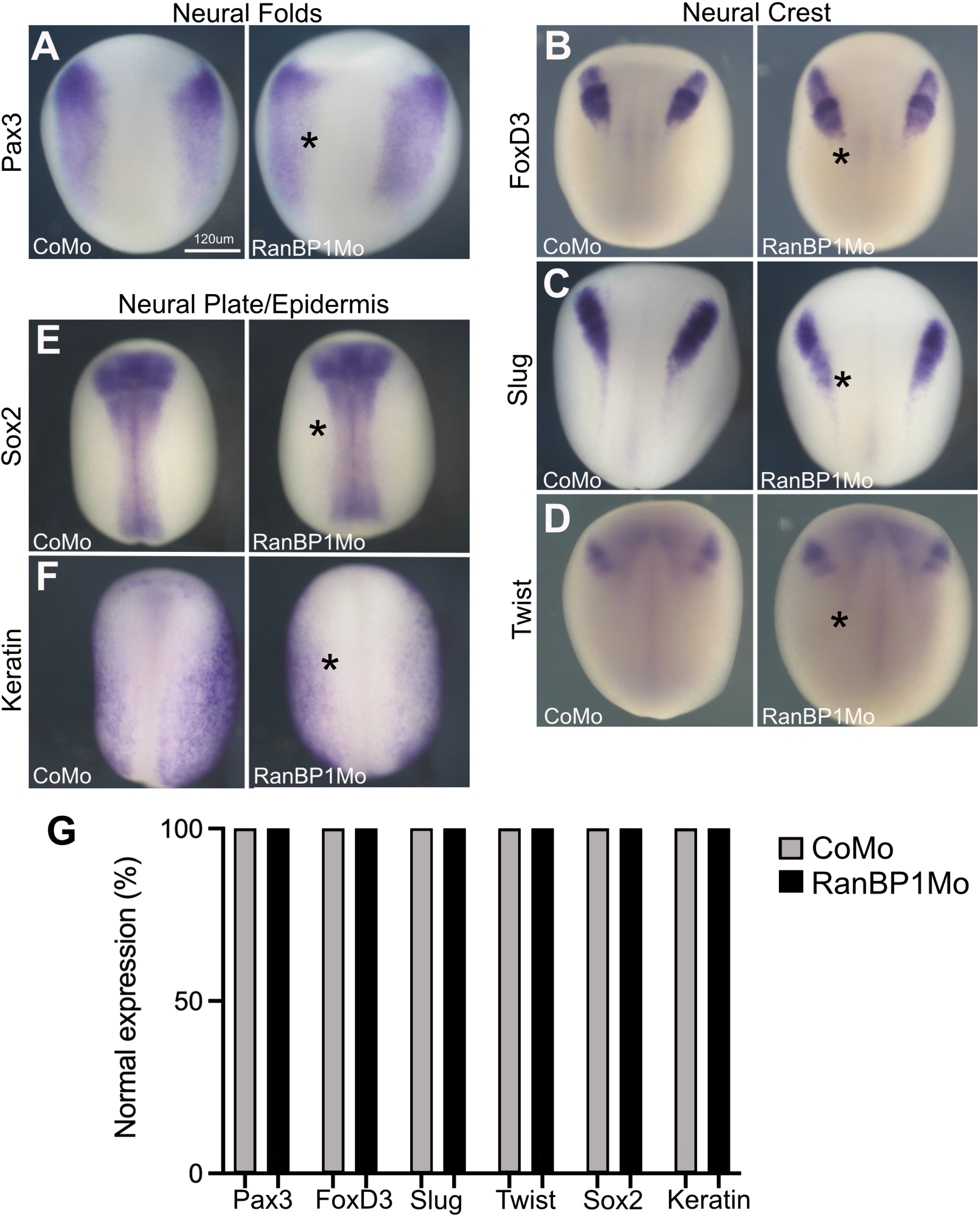
RanBP1 depletion does not affect early NC or ectodermal specification. 2-cell stage embryos were injected with a Control MO or RanBP1 MO in one of the two blastomeres. Embryos were fixed at stage 14 (**A**), 16 (**B-D**) and 18 (**E,F**), and the indicated genes were analyzed by ISH. The injected side is indicated by an asterisk. No effect on the expression of the markers was observed upon RanBP1 MO injection. (**G**) Quantification showing the % of embryos with normal expression. At least 15 embryos were analyzed for each gene/condition.

**Figure 4.**
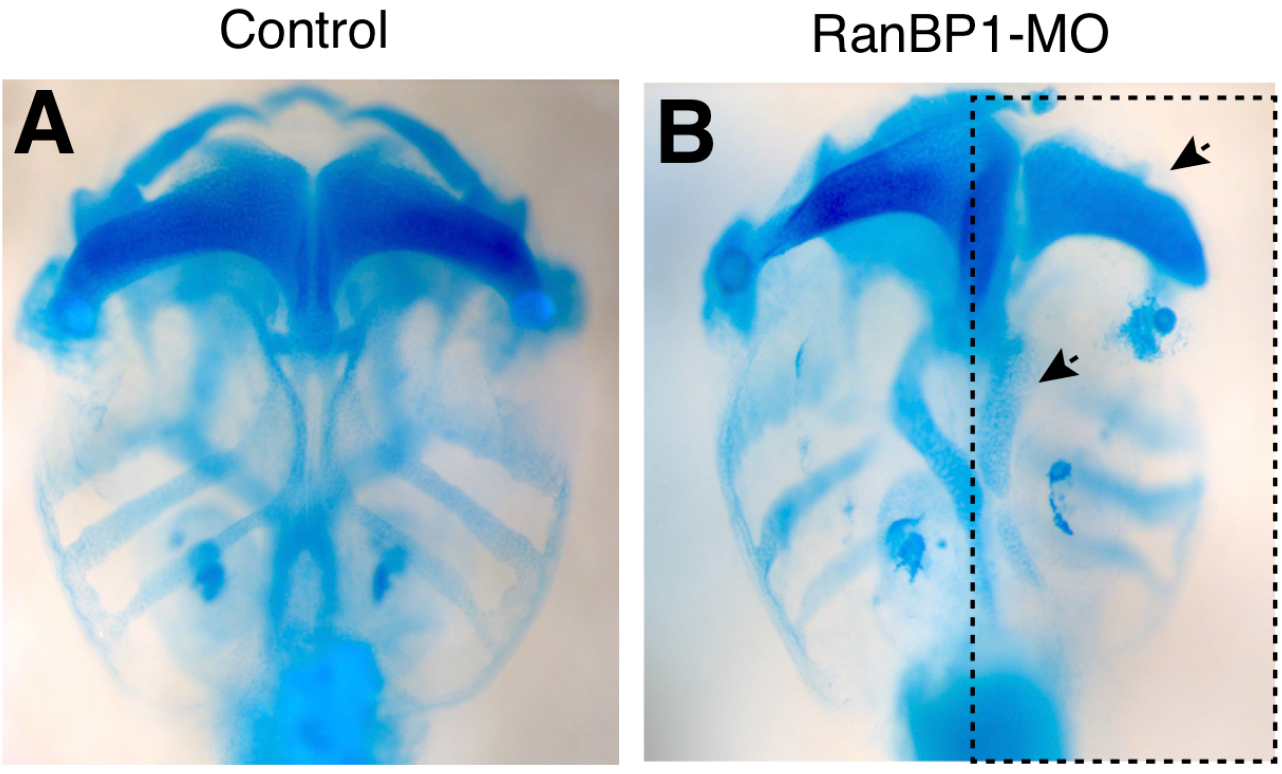
Decreasing RanBP1 expression leads to skeleton defects. (**A-B**) Cartilage labelled using Alcian Blue in control and RanBP1MO embryos. (**A**) is a wild type embryo and (**B**) one treated using the RanBP1 MO. At least 10 embryos were analysed for each genotype, and the picture presented in (**A-B**) are representative of the RanBP1 MO phenotype.

### RanBPl is required for NC chemotaxis in response to SDF-l exposure

Next, to examine the possible cause for the NC migration phenotype we observed in RanBP1-MO embryos, we turned to an *ex vivo* chemotaxis assay, making use of beads coated with Stromal Derived Factor 1 SDF-1(Theveneau et al., 2010). In this assay, the beads are fixed into a fibronectin coated dish and NC migration can be recorder using time-lapse microscopy (Figure 5A,B and Supplementary Movie S1). Wild type NC migrate towards the SDF-1 bead as a collective showing high persistent directionality (Figure 5B). In contrast, RanBP1-MO injected cells failed to chemotax towards SDF-1 (Figure 5B).

**Figure 5:**
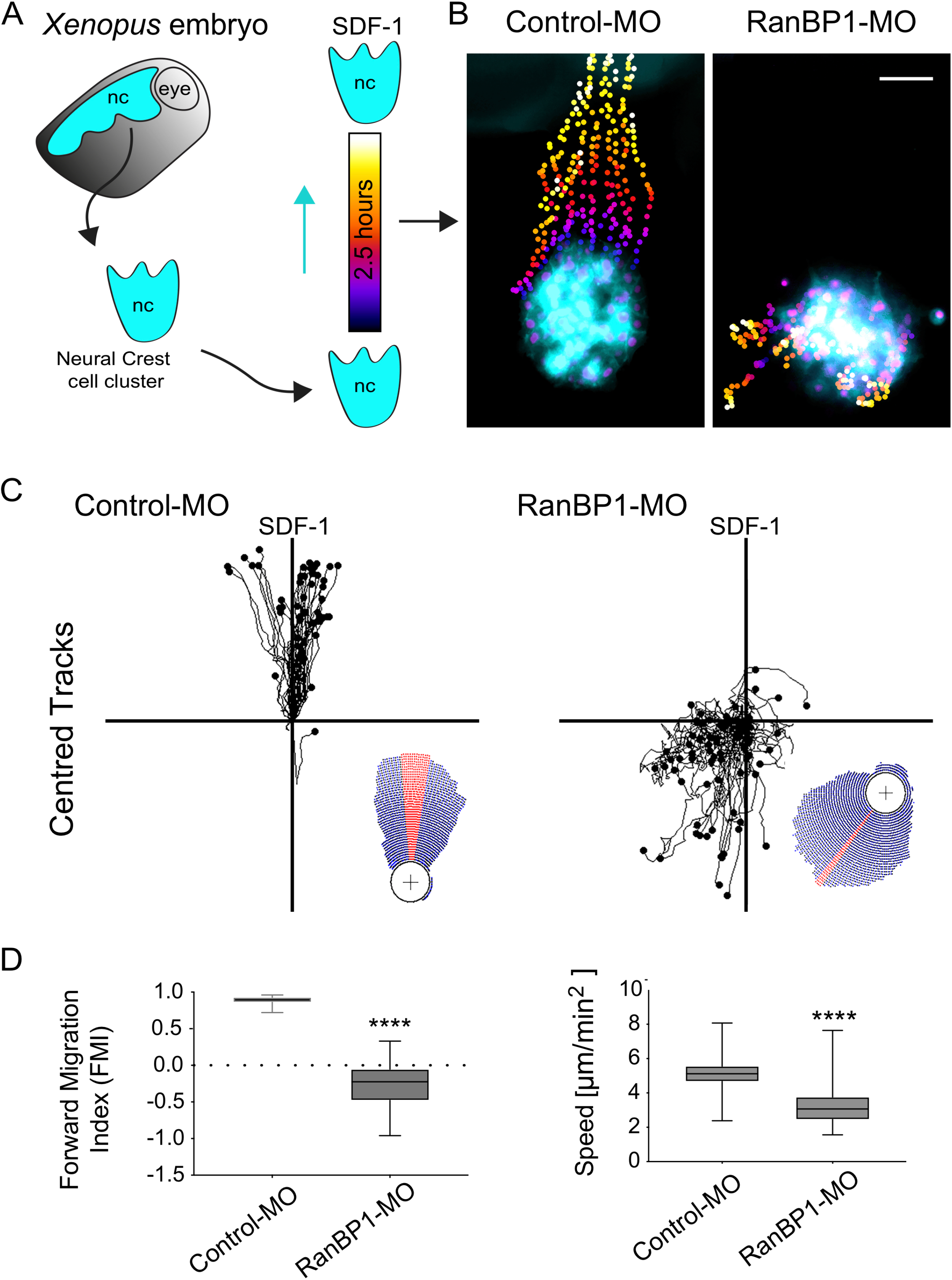
RanBP1 is required for NC chemotaxis towards SDF-1. (**A**) Diagram depicting our chemotaxis assay. (**B**) Clusters of Control or RanBP1-MO injected NCs where their trajectories have been colour coded as a function of time. The Scale bar represents 50 μm. (**C**) Centred tracks showing the trajectories of multiple Control or RanBP1 treated NCs. (**D**) Box plot showing the FMI and speed of migration of Control and RanBP1-MO cells, bars represent median and wiskers the min and max values; two-tailed Mann–Whitney test ***, P < 0.001; CI= 95%; ncontrol= 43, nRanBP1-MO= 68 cells from three independent experiments.

In order to gain further insight into this phenotype, we tracked the migrating NC to extract their trajectories and then calculated how persistent their migration towards SDF-1 was, by determining their forward migration index. A forward migration index close to 1 indicates sustained directional cell migration. While control NC efficiently chemotax with fairly straight trajectories and a forward migration index of ~ 1, we found that RanBP1-MO injected cells displayed more randomised trajectories with an average forward migration index of ~0.4; a typical value displayed by clusters that fail to show directionality during migration (Foxman et al., 1999) (Figure 5C and Supplementary Movie S1). From this, we conclude that RanBP1 is required for directed migration of NC, in response to SDF-1.

### RanBP1 is required for directed migration

Failure in chemotaxis could be due to many reasons, including a failure for individual cells to polarize along the front-to-rear axis. In order to examine this possibility, we tracked individual NC to extract cell motility parameters including migratory speed, directionality and circularity (Figure 6A-D and Supplementary Movie 2 and 3). Consistent, with our previous observations, quantification of RanBP1-MO individual NCs showed that these cells display slower migratory speed and less directionalities when compared to wild type cells (Figure 6B-C; Supplementary Movie 2 and 3). In addition, many cells failed to extend migratory protrusions and displayed a rounder morphology than that of wild type cells, which is indicative of poor front-to-rear polarity and defect in NC migration (Figure 5D and Supplementary Movie 2 and 3). Altogether these results indicate that RanBP1 is required in NC to support directed migration, in response to SDF-1.

**Figure 6:**
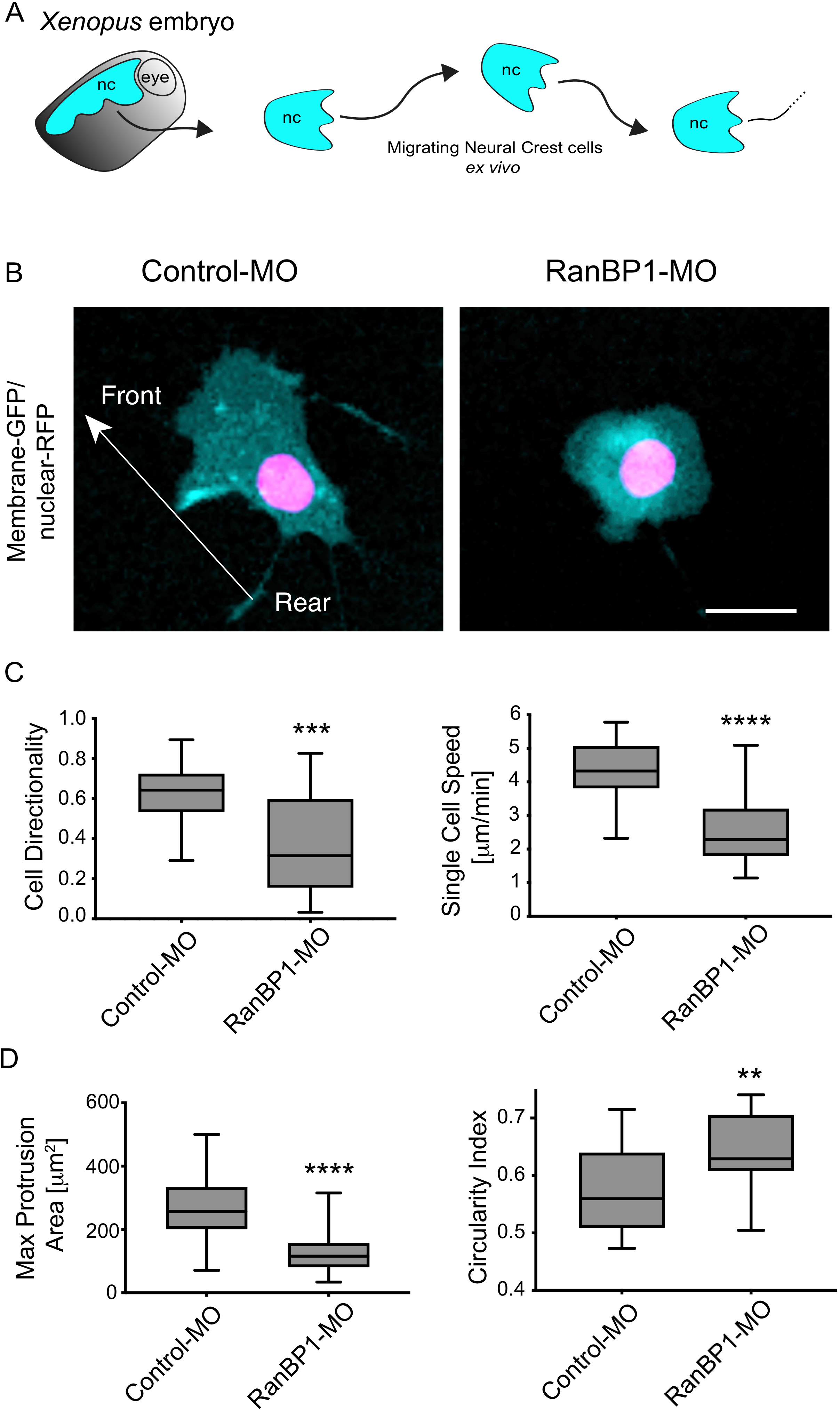
RanBP1 is required for NC polarity and motility. (**A**) Drawing showing our single NCs migration assay. (**B**) Representative images of migrating Control-MO or RanBP1-MO NCs clusters, showing loss of rear-front polarity. (**C-D**) Parameters for individual cell motility as obtained from Supplementary Movie 3. Box plots showing cell directionality and speed of Control and RanBP1-MO cells in (**C**) and max protrusion area as well as circulatiry index in (**D)**. In (**C-D**) bars represent median and wiskers the min and max values. In (**C**), for single cell directionality two-tailed Mann–Whitney test ***, P < 0.0005; ncontrol= 16, nRanBP1-MO= 36 cells from three independent experiments; and for single cell speed two-tailed t-test ****, P < 0.0001; CI= 95%; ncontrol= 20, nRanBP1-MO= 34 cells from three independent experiments. In **(D)**, for max protrusion area two-tailed Mann-Whitney test ****, P < 0.0001; n_control_= 185, n_RanBP1-MO_= 191 protrusions from three independent experiments; and for circulatiry index two-tailed t-test **, P = 0.0062; CI= 95%; n_control_= 20, n_RanBP1-MO_= 20 cells from three independent experiments. Scale bar = 20 micron.

### LKB1 mediates RanBP1 function during neural crest cell migration

In cortical neurons, RanBP1 funtion in polarity is principaly mediated by the kinase LKB1 (Mencarelli et al., 2018). LKB1 is expressed and required in NC cells for their migration (Creuzet et al., 2016), raising the issue that the RanBP1-LKB1 pathway is at play during this process. To test this idea, we took advantage of a previously published *Xenopus* LKB1 kinase dead dominant negative construct (Ossipova et al., 2003). We found that the injection of this construct leads to a failure in NC migration that resembles that of RanBP1 injected animals (Figure 6A-C). Then, to determine whether LKB1 could mediate part of RanBP1 function in NC migration, we performed an epistatic experiment whereby RanBP1-MO was co-injected with a wildtype version of LKB1. This co-injection of *LKB1* mRNA was sufficient to restore the defects observed upon knocking down RanBP1 expression (Figure 7D-F). These results are consistent with LKB1and RanBP1 functioning together to enable directed NC migration.

**Figure 7:**
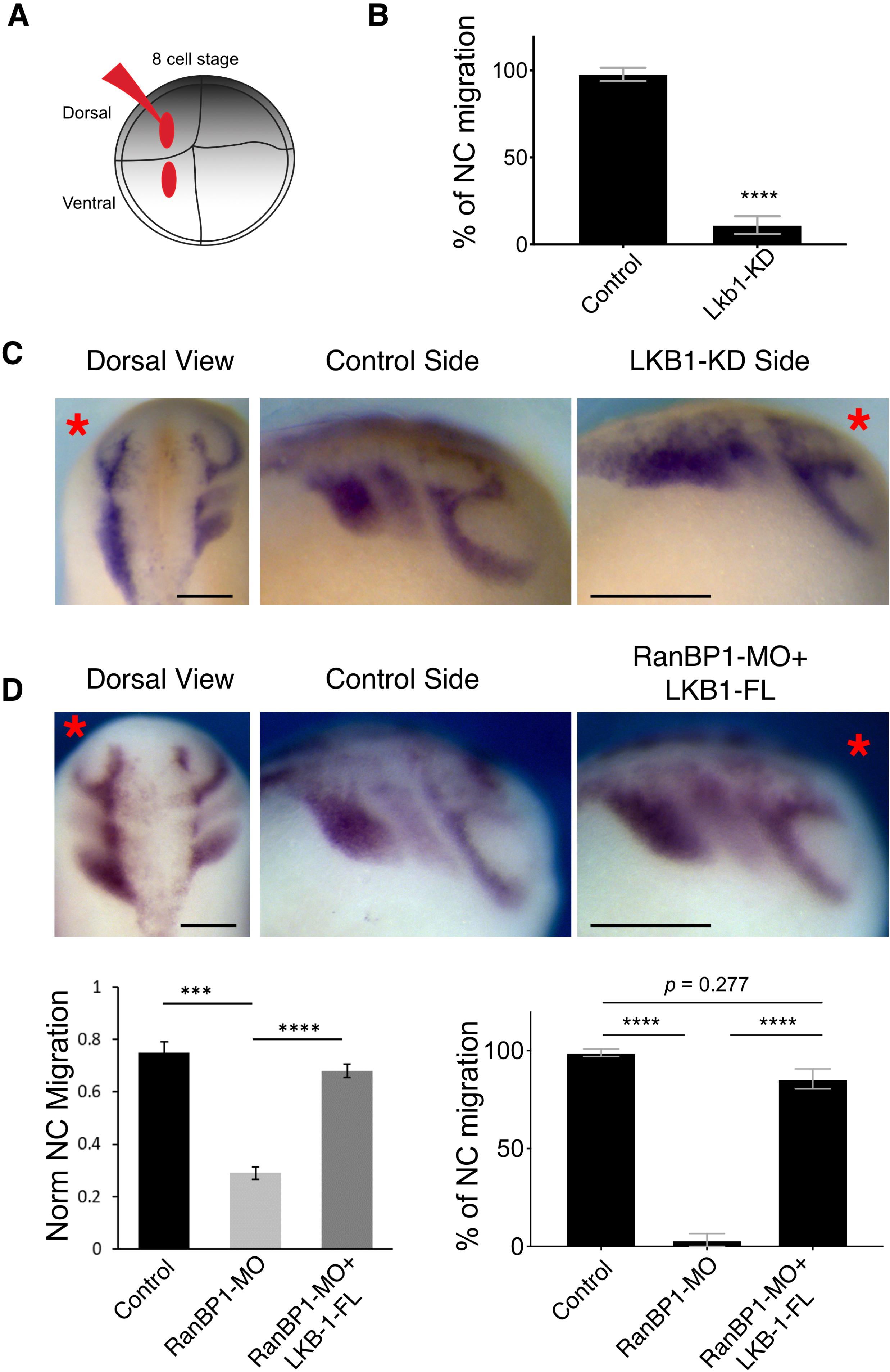
LKB1 expression can rescue the RanBP1-MO migration phenotype. (**A**) Diagram showing our injection strategy and (**B**) histogram representing the percentage of Control or LKB1-KO injected embryos that displayed NC migration defects. (**C-D**) Dorsal and lateral views of embryos hybridized with a probe against *snail2* to label migrating NCs. (**C**) A red asterisk marks the injected side of the embryo. Lateral views show that while NC form the control side of the embryo migrate normally by forming dorso ventral streams, LKB1-KD cells (red asterisk) remain in the dorsal part of the embryo. (**A**) A red asterisk denotes the injected side of the embryo and dorsal as well as lateral views show that both Control and RanBP1-MO+LKB1mRNA co-injected NCs normally migrate by forming dorso-ventral streams. Scale bars represent 200 μm. (**E**) Normalized NC migration and percentage of embryos displaying the representative phenotypes shown here. Histograms represent the media and bars the standard deviation of three independent experiments, N= 20 embryos for each condition. two-tailed t-test ***, P < 0.001 and ****, P < 0.0001.

## DISCUSSION

Nuclear export of LKB1 is regulated by Ran and its effector RanBP1 (Baas et al., 2003; Dorfman and Macara, 2008), and is required for polarization of epithelial cells and rat cortical neurons (Baas et al., 2004; Matsuki et al., 2010; Mencarelli et al., 2018; Shelly et al., 2007). Here we show that RanBP1 and LKB1 are also required for directed migration of cephalic NCs. In this context, our results indicate RanBP1 is required for NCs to polarize properly along the front-to-rear axis, and thus for chemotaxis in response to a source of SDF-1. Altogether, our work is compatible with a model whereby RanBP1 requirement in NC migration stems from its function in promoting unloading of LKB1 from the nuclear export complex, essential for releasing this kinase in the cytosol.

### RanBP1 is required for NC front-rear polarization

RanBP1 is required in culture cortical neurons for axon formation and *in vivo,* for neuron migration duing cortex development (Mencarelli et al., 2018). As it is the case in cortical neurons, RanBP1 is essential for cell viability in NC cells (Data not shown). However, we were able to calibrate our MO concentration to achieve an hypomorphic situation, whereby the expression of this factor is not totally inhibited. In this hypomorphic situation, cell viability is not affected, but we expect that RanBP1-dependent cargo release is less efficient compared to wildtype cells. In this case, we find a clear requirement for RanBP1 in front-rear polarity in individual migrating NC. RanBP1 has been associated with velo-cardio-facial/DiGeorge syndrome {Dunham, 1999 #3338;Merscher, 2001 #3339, whch includes phenotypes that can be linked to NC biology. Our results thefore raise the possibility that RanBP1 deficiency contribute to this syndrome by affecting NC migration.

### The RanBP1-LKB1 axis in NC polarity

Collective NC migration has been shown to be triggered by stiffening of their migratory substrate {Barriga, 2018 #3327}. As they start migrating, these cells need to integrate several extrinsic cues to orient their polarity and to move towards their target tissues (Barriga and Theveneau, 2020; Shellard and Mayor, 2020). One of the most studied mechanisms of NCs guidance is their chemotaxis in response the chemokine SDF-1, which is mediated by the CCR4 receptor and the small GTPase Rac1 (Theveneau et al., 2010). However, how CCR4 activation translates into polarization of NC cell is not well understood. LKB1 has also been proposed to promote NC migration through AMP-activated protein kinase and via actomyosin regulation through the Rho-dependent kinase ROCK (Creuzet et al., 2016). Our work confirms LKB1 is required for NC cell migration, and links this key polarity factor to the nucleocytoplasmic transport pathway, through RanBP1.

LKB1 has been shown to regulate polarity in a range of cell types and systems. This kinase is essential for enabling asymmetric division of the C. elegans zygote (Watts et al., 2000). It has also been shown to promote neuron polarity and dendrite specification by regulating microtubule stability through the family of MARK-kinases and in particular through the related SAD-A/B kinases, and through Golgi regulation (Barnes et al., 2007; Huang et al., 2014; Matsuki et al., 2010; Mencarelli et al., 2018; Shelly et al., 2007). Morevover, LKB1 regulates apical-basal polarity in epithelial cells (Amin et al., 2009; Baas et al., 2004; Martin and St Johnston, 2003). In neurons and human epithelial cells, LKB1 function in polarity starts with its nuclear export (Baas et al., 2004; Mencarelli et al., 2018). Our results show that overexpression of LKB1 can rescue the RanBP1 loss-of-function phenotype. While we cannot rule out that LKB1 functions in paralelle to RanBP1 in NC migration, and that overexpression of this kinase can compensate for the inhibition of RanBP1 expression, we favor a model whereby RanBP1 function in NC migration is to control the LKB1 availabilty in the cytosol of the migrating NC cells. In this model, response to SDF-1 stimulation requires LKB1.

---

**Supplementary Video 1. RanBP1 is required for NC chemotaxis towards SDF-1.**

*ex-vivo* time-lapse of control and RanBP1-MO NC migrating in an SDF-1 chemotaxis assays. Time-lapse setting was set at 1 picture every 5 min; 35 frames are shown. Frame rate is 7 frames per second.

**Supplementary Video 2. RanBP1 is required for cluster cell spreading.**

*ex-vivo* time-lapse of control and RanBP1-MO NC migrating in a cluster spreading assay. Time-lapse setting was set at 1 picture every 5 min; 54 frames are shown.

Frame rate is 7 frames per second.

**Supplementary Video 3. RanBP1 is required for single cell motility.**

*ex-vivo* time-lapse of control and RanBP1-MO NC migrating in a single cell migration assay. Time-lapse setting was set at 1 picture every 5 min; 31 frames are shown. Frame rate is 7 frames per second.

## EXPERIMENTAL PROCEDURES

### *Xenopus* manipulation

Adult *X. laevis* were maintained in a range of temperatures 14-18 °C. To obtain embryos we followed established protocols (Barriga et al., 2013). Briefly, superovulation was induced in adult female frogs by injecting chorionic gonadotrophin (Intervet), then obtained oocytes were mixed with a sperm solution for *in vitro* fertilizations. Embryos were staged as previously described (Hubrecht-Laboratorium (Embryologisch Institut), 1967) and maintained at 14°C. All animal experiments were approved by the Biological Service Unit at University College London and complied with UK Home Office guidelines (Animal Act 1986); or by the Ethics Committee and the Animal Welfare Body of the Instituto Gulbenkian de Ciencia and by the Direção Geral de Alimentação e Veterinária (DGAV), Portugal.

### *In situ* hybridisation and *in vitro* transcription

*In situ* hybridizations and templates for all *in vitro* transcription were generated as previously described (Barriga et al., 2013; Barriga et al., 2019; Theveneau and Mayor, 2011). Digoxigenin-labeled RNA probes against *snail2* (Mayor et al., 1995), *twist* (Hopwood et al., 1989), *FoxD3* (Kelsh et al., 2000), *Sox2* (Kishi et al., 2000), *Keratin* (Jonas et al., 1985), *Pax3* (Alkobtawi et al., 2018) was used in this study. In brief, digoxigenin-labeled probes against target mRNA were transcribed by using an *in vitro* Transcription System (Promega P1420) and all mRNAs were transcribed with a mMESSAGE mMACHINE SP6 and T7 Transcription Kits (Thermo-Fisher AM1340 and AM1344, respectively).

#### Analysis of neural crest migration *in vivo*

After imaging embryos subjected to *in situ* hybridization, the dorso-ventral length of the NC was extracted with the built-in measurement tool of ImageJ. This length was normalized against the total dorso-ventral length of the embryo that was also extracted using ImageJ. Further analyses and comparisons were carried out as described in the Statistical analysis section.

### Morpholino and mRNA injections

Injections were performed using glass needles calibrated to deliver 10nL. Briefly, at eight-cell stage a ventral and dorsal animal blastomeres were injected with: 5ng of a morpholino designed against *Xenopus RanBP1 (RanBP1-MO* 5’-GCGGGTCTTCTGTTTCTACAATGTC-3’), with the same amount of a standard control MO and/or with 1ng of mRNAs encoding for Lkb1 or RanBP1. Similarly, to fluorescently tag NCs, 250 pg of membrane GFP (mGFP) and or nuclear RFP (nRFP) were injected, as shown in each figure.

### Neural crests dissection

Dissections were performed after (Barriga et al., 2019). In brief, the vitellin membrane was mechanically removed, and embryos were immobilized in a dish containing modelling clay. Then, the epidermis was removed with a hair knife and the same tool was used to explant the NCs. NCs explants were maintained in Danilchik’s medium (DFA; 1 mM MgSO4(7H2O), 5mM Na2CO3, 4.5 mM KGluconate, 53 mM NaCl, 32 mM NaGluconate, 0.1% BSA and 1 mM CaCl2; after dissolving the components the pH was adjusted to 8.3 with Bicine).

### Single cell migration assay

To analyse motility at the single cell level, we used an assay in which explants were mechanically disaggregated to obtain single cells, and then platted onnto a dish coated with fibronectin (μ-Dish, 35 mm, Ibidi). NCs migration was recorded by time-lapse microscopy.

### Chemotaxis assay

NC response to SDF-1 was assesed after(Theveneau and Mayor, 2011). In brief, acrylic heparin beads (Sigma-Aldrich) were coated with 1 μg/ml Sdf-1 (Sigma-Aldrich) for 1 hour, washed with PBS. Silicon grease was used to fix these SDF-1-coated beads to a fibronectin-coated dish (Ibidi). NCs clusters (tagged with mRFP and or mGFP) were placed in front of the bead and their directional response was recorded by time-lapse.

### Time-lapse imaging

For chemotaxis and single cell migration assays, an emage was acquired every 5 min at 18 °C using an upright microscope (DMR XA2, Leica). This microscope was equipped with a motorized stage (Prior Scientific) to facilitate imaging of multiple samples and a high resolution camera (Orca-5G Hamamatsu). Images acquisition was performed with a 20× objective (HCX APO L 20×/0.50 W U-V-I FWD = 3.50 mm, Leica) and all microscopes components were motorised and controlled through a dedicated software, SimplePCI (Hamamatsu).

### *In situ* hybridization imaging

Embryo images were acquired by using 3.2× magnification using a Leica MZ FL III dissecting microscope equipped with a Leica DFL420 camera, controlled through a Leica IM50 imaging software.

### Cell tracking, chemotaxis, protrusion area and circularity index

The trajectories of individual cells or clusters were established using a standard Manual tracking plugin for ImageJ and the Chemotaxis Tool plugin from the same software was used to extract speeds of migration and directionality. Max protrusion area and circularity index were extracted by using the built-in hand free selection and measurement tools of ImageJ. Data were further analysed as described in Statistical analysis.

### Image processing

The ImageJ software was used to generate *z*-stacks, maximum projections and to mount timelapse movies. ImageJ and/or Adobe Photoshop was used for linear adjustment of display map levels, crop for re-sizing and application of pseudocolour. Adobe Illustrator was used to assemble all figures.

### Statistical analysis

Each experiment was repeated at least three times. To determine whether data set were parametric or not we used Kolmogorov–Smirnov, d’Agostino–Pearson or Shapiro–Wilk test in Prism7 (GraphPad). When parametric, significance was compared by using unpaired *t*-test (two-tailed, unequal variances) and in case of non-parametric distribution, significance between ranks was determined with Mann–Whitney *U*-test (two-tailed). CI was set at 95% in all cases.

## ACKNOWELDGEMENTS

We are grateful to the Pichaud lab for discussions related to this study, and the LMCB, CDB and IGC imaging, and the aquatic animal facility at UCL and IGC for their support. The authors also thank Prof Jeremy Green for kindly providing the *Xenopus* LKB1 constructs. This work was funded by an MRC (MC_UU_12018/3), BBSRC (BB/R000697) and Royal Society grant (Award #181274) to F. Pichaud, and benefited from MRC core funding to the LMCB, covering access to microscopy (MC_12266B). E.H. Bariga was funded at UCL through a BBSRC grant (Grant No: M008517) to R. Mayor. Work in R. Mayor lab is funded by grants from the Medical Research Council (MR/S007792/1) and BBSRC (M008517). Work in EHB lab is funded by the European Research Council (ERC) (grant agreement No. 950254), EMBO IG Project Number 4765; la Caixa Junior Leader Incoming (94978) and support provided by Instituto Gulbenkian de Ciencia and Fundaçao Calouste Gulbenkian (start-up fund I-411133.01).

## Notes

### Competing Interest Statement

The authors have declared no competing interest.

